# Paxlovid-like nirmatrelvir/ritonavir fails to block SARS-CoV-2 transmission in ferrets

**DOI:** 10.1101/2022.11.20.517271

**Authors:** Robert M Cox, Carolin M Lieber, Josef D Wolf, Amirhossein Karimi, Nicole A P Lieberman, Zachary M Sticher, Pavitra Roychoudhury, Meghan K Andrews, Rebecca E Krueger, Michael G Natchus, George R Painter, Alexander A Kolykhalov, Alexander L Greninger, Richard K Plemper

## Abstract

Despite the continued spread of SARS-CoV-2 and emergence of variants of concern (VOC) that are capable of escaping preexisting immunity, therapeutic options are underutilized. In addition to preventing severe disease in high-risk patients, antivirals may contribute to interrupting transmission chains. The FDA has granted emergency use authorizations for two oral drugs, molnupiravir and paxlovid. Initial clinical trials suggested an efficacy advantage of paxlovid, giving it a standard-of-care-like status in the United States. However, recent retrospective clinical studies suggested a more comparable efficacy of both drugs in preventing complicated disease and case-fatalities in older adults. For a direct efficacy comparison under controlled conditions, we assessed potency of both drugs against SARS-CoV-2 in two relevant animal models; the Roborovski dwarf hamster model for severe COVID-19 in high-risk patients and the ferret model of upper respiratory tract disease and transmission. After infection of dwarf hamsters with VOC omicron, paxlovid and molnupiravir were efficacious in mitigating severe disease and preventing death. However, a pharmacokinetics-confirmed human equivalent dose of paxlovid did not significantly reduce shed SARS-CoV-2 titers in ferrets and failed to block virus transmission to untreated direct-contact ferrets, whereas transmission was fully suppressed in a group of animals treated with a human-equivalent dose of molnupiravir. Prophylactic administration of molnupiravir to uninfected ferrets in direct contact with infected animals blocked productive SARS-CoV-2 transmission, whereas all contacts treated with prophylactic paxlovid became infected. These data confirm retrospective reports of similar therapeutic benefit of both drugs for older adults, and reveal that treatment with molnupiravir, but not paxlovid, may be suitable to reduce the risk of SARS-CoV-2 transmission.

## Introduction

Vaccines and antivirals have helped to limit disease severity in the coronavirus disease 2019 (COVID-19) pandemic. However, pandemic viral spread continues, affecting even those with vaccine or naturally-acquired immunity^1-3^. The rise of new SARS-CoV-2 variants of concern (VOC) capable of escaping preexisting immunity has undermined the hope of rapidly ending the pandemic through large-scale vaccination campaigns^1,4,5^, and indirect immune imprinting may never allow vaccine boosters to be successful against future viral evolution. In 2021, the FDA granted emergency use authorizations for two orally available antivirals, molnupiravir and paxlovid^6,7^. Molnupiravir, a prodrug of the broad-spectrum nucleoside analog N^4^-hydroxycytidine^8^, targets the viral RNA polymerase, triggering lethal viral mutagenesis^9,10^. Paxlovid consists of two components, the peptidomimetic viral main protease (M^pro^) inhibitor nirmatrelvir and the cytochrome P450 enzyme inhibitor ritonavir^11^, which is required to improve the oral availability of nirmatrelvir. Having formed covalent dead-end complexes with nirmatrelvir, M^pro^ is unable to process the viral polyprotein, interrupting the viral replication cycle. Initial phase II/III clinical trials suggested superior efficacy of paxlovid compared to molnupiravir, claiming 89% versus 30% reduction in patient hospitalization and death^12,13^.

These initial data and extensive public discussion of possible carcinogenic potential of molnupiravir should it be incorporated into host DNA triggered elevation of paxlovid to standard-of-care (SOC)-like status for treatment of vulnerable patient populations in the United States^14^. However, a recently completed carcinogenicity study in the Tg.rasH2 mouse model to assess carcinogenic potential of molnupiravir demonstrated that continuous dosing for 6 months was not carcinogenic^15^, addressing speculations about drug safety. Recent large-scale clinical trials and retrospective assessments of therapeutic benefit have revealed that efficacy of paxlovid is substantially lower than initially seen against VOC delta^12,16,17^, whereas therapeutic benefit of molnupiravir in older adults was underestimated^15,17^. In these studies, neither drug provided significant therapeutic benefit to younger adults below 65 years of age^12,16,18^.

Assessing the effect of post-exposure prophylactic treatment of household contacts of a confirmed SARS-CoV-2 case with paxlovid, the EPIC-PEP trial revealed no significant effect of paxlovid drug on preventing transmission^19,20^. Clinical reports demonstrated rebounding virus replication in patients who have received paxlovid^2,21,22^ and anecdotal cases of SARS-CoV-2 transmission at rebound have further questioned whether paxlovid may provide epidemiologic benefit through reducing the risk of SARS-CoV-2 spread. Using the ferret SARS-CoV-2 transmission model, we have established in previous work that oral molnupiravir blocks direct-contact transmission of the virus to untreated sentinels within 18 hours of treatment start^23,24^.

In this study, we employed two relevant animal models to evaluate paxlovid versus molnupiravir benefit under experimentally controlled conditions; the Roborovski dwarf hamster model for acute respiratory failure in highly susceptible patients with elevated risk of lethal outcome^24,25^ and the ferret model for upper respiratory tract disease and transmission^23,26,27^. Both drugs met the primary efficacy endpoint of ensuring complete survival of dwarf hamsters infected with VOC delta or omicron and effect size of lung titer reduction was comparable. However, ferrets receiving a human-equivalent dose of paxlovid continued to shed virus into nasal lavages, had virus replication rebound after two days of treatment, and transmitted the virus efficiently to all sentinels, whereas molnupiravir fully suppressed SARS-CoV-2 transmission. Prophylactic treatment of sentinels with molnupiravir, but not paxlovid, prevented infection by untreated close-contacts.

## Results

To validate sourced nirmatrelvir stocks (extended data Fig. 1a), we determined antiviral potency on cultured VeroE6-TMPRSS2 cells against SARS-CoV-2 isolates representing lineage A (USA-WA1/2020), lineage B.1.1.7 (VOC alpha; hCoV-19/USA/CA/UCSD_5574/2020), lineage B.1.351 (VOC beta; hCoV-19/South Africa/KRISP-K005325/2020), lineage P.1 (VOC gamma; hCoV-19/Japan/TY7-503/2021), lineage B.1.617.2 (VOC delta; clinical isolate #233067), lineage B.1.1.529 (VOC omicron; hCoV-19/USA/WA-UW-21120120771/2021) lineage BA.2 (clinical isolate 22012361822A), lineage BA.2.12.1 (hCoV-19/USA/WA-CDC-UW22050170242/2022), and linage BA.4 (hCoV-19/USA/WA-CDC-UW22051283052/2022) (extended data Fig. 1b).

### *In vitro* efficacy of nirmatrelvir against current VOC

Infected cells were incubated with serial dilutions of nirmatrelvir in media containing 2 µM CP-100356, a P-glycoprotein inhibitor added to limit rapid compound efflux^28^. Nirmatrelvir exhibited nanomolar potency against all VOCs tested, returning half-maximal (EC_50_) and 90% maximal (EC_90_) concentrations ranging from 21.1 to 327.6 nM and 32.8 to 421.4 nM, respectively, which recapitulated results of previous *in vitro* dose-response studies^29,30^.

### Prevention of severe SARS-CoV-2 disease in Roborovski dwarf hamsters

To identify a drug dose that results in nirmatrelvir plasma exposure in Roborovski dwarf hamsters at least equivalent to that seen in humans, we determined single-oral dose PK properties of nirmatrelvir delivered alone or in combination with ritonavir. Hamsters were administered 250 mg/kg nirmatrelvir with or without 83.3 mg/kg ritonavir. In the absence of ritonavir, 250 mg/kg nirmatrelvir delivered plasma exposure levels below those achieved in humans taking paxlovid (Fig. 1a, extended data Table 1). When combined with ritonavir, we noted nirmatrelvir plasma exposure exceeding that observed in humans by 7.6-fold^31,32^, confirming use of a conservative human equivalent dose (HED) of paxlovid.

**Figure 1.**
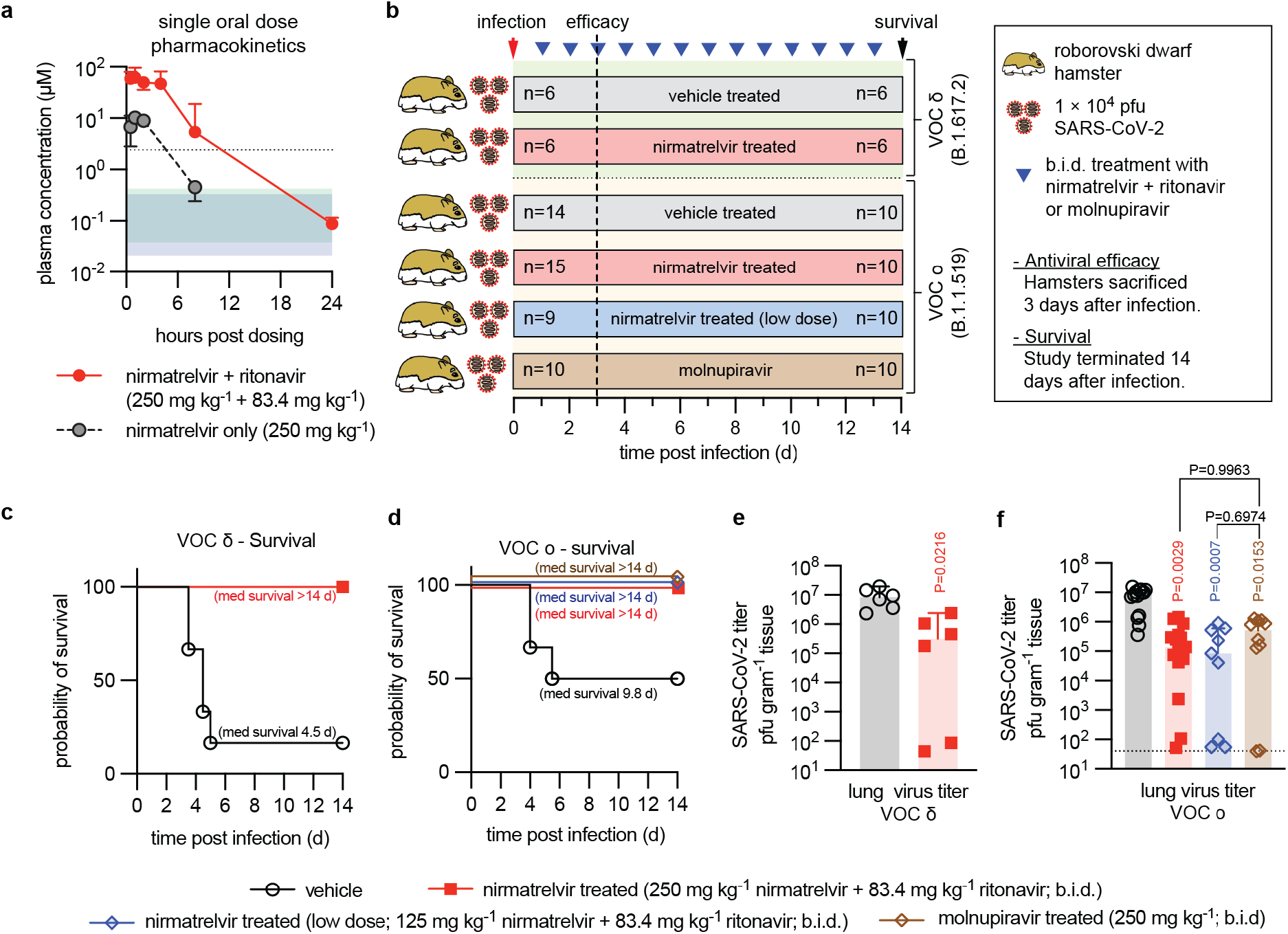
Efficacy of nirmatrelvir in Roborovski dwarf hamsters. **a**, Plasma concentration of nirmatrelvir over 24-hours after dosing. Single oral dose of nirmatrelvir (250 mg/kg), with (red circles) or without (black circles) ritonavir (83.4 mg/kg). Symbols show group medians. **b**, Antiviral efficacy study schematic. Animals were infected intranasally with 1×10^4^ pfu of VOC delta or omicron and monitored for 14 days after infection. Clinical signs were assessed once daily (blue triangles). Viral load was determined in a duplicate set of animals 3 days after infection. **c-d**, Survival curves of VOC delta (c) or omicron (d) infected dwarf hamsters from (b) Log-rank (Mantel-Cox) test, median survival (med survival) is shown. **e-f**, Lung virus titers 3 days after infection with VOC delta (e) or omicron (f) as shown in (b). Symbols represent independent biological repeats, columns show group medians with 95% CIs. Significance was determined using unpaired 2-tailed t tests (e) or 1-way ANOVA with Tukey’s post-hoc test (f); P values are shown.

Matching the design of our previous molnupiravir efficacy studies in the dwarf hamster model^24^, we initiated oral treatment at this dose level 12 hours after intranasal infection of the dwarf hamsters with 1 × 10^4^ pfu of VOC delta or omicron, and continued treatment in a b.i.d. regimen (Fig. 1b). We included an additional low dose group (125 mg/kg nirmatrelvir and 83.3 mg/kg ritonavir) for VOC omicron to assess dose-dependency of potency and effect size. The 12-hour time to treatment onset reflected rapid disease progression in the dwarf hamster model^24^, which mimics human patients at transition stage to complicated COVID-19. As we have reported for molnupiravir treatment^24^, all treated dwarf hamsters met the primary efficacy endpoint and survived infection independent of VOC used (Fig. 1c-d). By contrast, 83% and 40% of vehicle-treated animals had succumbed to the disease 6 days after infection with VOC delta and omicron, respectively. Drug treated animals also displayed minimal to no clinical signs, whereas dwarf hamsters in vehicle groups experienced hypothermia and body weight loss (extended data Fig. 2).

Virus burden in lung and trachea of duplicate sets of equally infected and treated dwarf hamsters was determined 3 days after infection to determine effect size of treatment. As we have reported previously, both VOC delta and omicron replicated efficiently in the dwarf hamsters, each reaching peak lung virus loads of approximately 10^7^ pfu/g lung tissue (Fig. 1e-f). Treatment of VOC delta-infected dwarf hamsters with paxlovid revealed a significant reduction in virus burden, resembling the reduction in VOC delta lung titer that we previously observed after treatment with molnupiravir^24^. However, effect size of treatment varied by several orders of magnitude between individual animals. We had noticed variation in effect size in molnupiravir-treated dwarf hamsters infected with VOC omicron before, but not VOC delta^24^. Head-to-head comparison of paxlovid and molnupiravir under identical experimental conditions in animals infected with VOC omicron demonstrated statistically significant reduction in lung virus load by either drug compared to vehicle-treated hamsters, but no statistically significant differences in effect size between both drugs (Fig.1f).

### Paxlovid treatment of SARS-CoV-2 infection in ferrets

To assess the impact of nirmatrelvir treatment on upper respiratory disease, we employed the SARS-CoV-2 ferret infection model^23,26,33,34^. To ensure cross-species consistency of nirmatrelvir PK performance, we again administered the compound first in a single oral dose at two levels, 20 mg/kg (predicted HED^31^) and 100 mg/kg (high-dose), each in combination with 6 mg/kg ritonavir. Overall nirmatrelvir exposure in ferrets administered HED was 32,700 hrs × ng/ml (Fig. 2a; extended data Table 2), which closely resembled that observed in humans of 28,200 hrs × ng/ml h^31,32^. Animals of the high-dose group reached nirmatrelvir plasma exposure levels of 49,500 hrs × ng/ml, indicating a dose-dependent, but not dose-proportional, PK profile.

**Figure 2.**
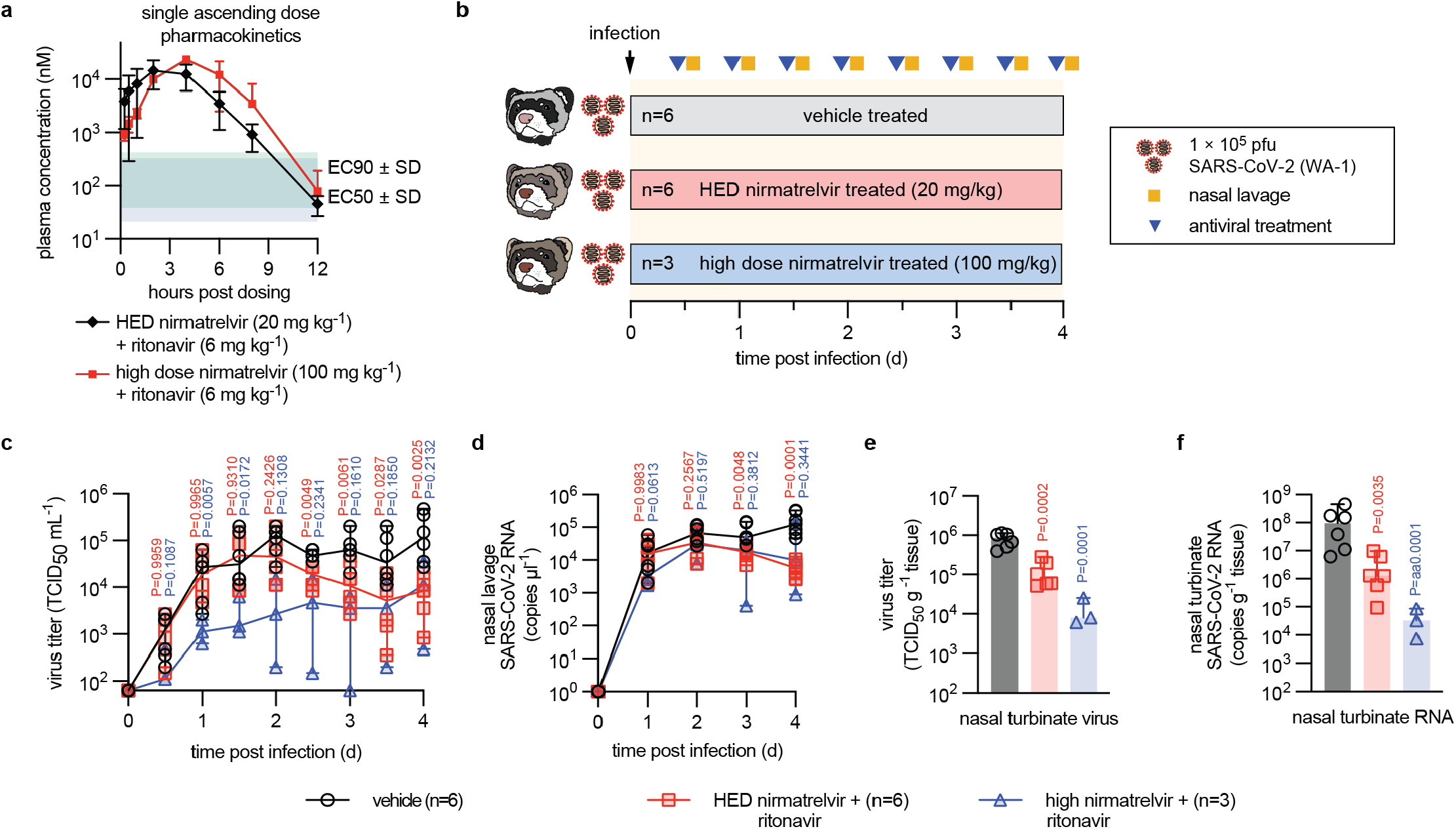
Efficacy of paxlovid in ferrets. **a**, Plasma concentration of nirmatrelvir determined over a 12-hour period after a single oral dose. Nirmatrelvir was dosed at 20 mg/kg (black diamonds, human equivalent dose (HED)) or 100 mg/kg (red squares, high-dose), each in combination with 6 mg/kg ritonavir. Symbols represent group means, lines intersect means, and error bars represent the standard deviation. **b**, Antiviral efficacy study schematic. Ferrets (n = 15 total) were infected intranasally with 1 × 10^5^ pfu of SARS-CoV-2 USA/WA-1. After 12 hours, groups of ferrets were treated orally with vehicle, HED, or high-dose paxlovid. **c-d**, Infectious SARS-CoV-2 titers (c) and SARS-CoV-2 RNA copies (d) in nasal lavages of animals shown in (b). **e-f**, Infectious titers (e) and SARS-CoV-2 RNA copies (f) in nasal turbinates extracted four days after infection. Symbols in (c-f) represent independent biological repeats (virus load of individual animals), lines intersect group medians (c-d), and columns (e-f) show group medians; error bars denote 95% CIs. Statistical analysis with 1-way (e-f) or 2-way (c-d) ANOVA with Dunnett’s (e-f) post-hoc tests; P values are shown.

Both dose levels were advanced to efficacy testing after intranasal infection of ferrets with 1 × 10^5^ pfu of SARS-CoV-2 isolate USA-WA1/2020 (lineage A), which most efficiently replicates and spreads in ferrets^23,24,33,35,36^ and other members of the Mustela genus^37,38^. Treatment was initiated 12 hours after infection and continued b.i.d., and nasal lavages collected twice daily to determine shed virus load (Fig. 3b). Characteristic for this model^23,26,33,34^, SARS-CoV-2-infected ferrets displayed no overt clinical signs (supplementary Fig. S1). Virus replicated efficiently in vehicle-treated animals, reaching plateau titers of approximately 10^5^ pfu shed virus and 10^5^ RNA copies per ml nasal lavage 48 hours after infection, which were maintained until study end on day 4 (Fig. 2c-d). Mean shed virus titers were lower in the HED group than vehicle control 36 hours after treatment onset, but these differences were not statistically significant. Ferrets receiving high-dose paxlovid experienced a significant titer reduction in nasal lavages, although even high-dose did not fully clear the infection, in contrast to treatment with molnupiravir at the species-adjusted human dose equivalent^39,40^. Rather, we noted a rebound of virus replication in both paxlovid dose groups 3.5 days after treatment start. Whole genome sequencing of virus populations in nasal lavage and turbinate samples taken from these animals revealed that most had acquired an N501T or Y453F substitution in the Spike protein that are characteristic for SARS-CoV-2 adaptation to replication in mustelids^23^ (extended data Fig. 3), but no signature resistance mutation to nirmatrelvir in nsp5^41-43^ emerged as allele dominant (uploaded as NCBI BioProject PRJNA894555). At study end, infectious titers and viral RNA copies in nasal turbinate tissues were statistically significantly reduced in paxlovid-treated animals compared to the vehicle group (Fig. 2e-f).

**Figure 3.**
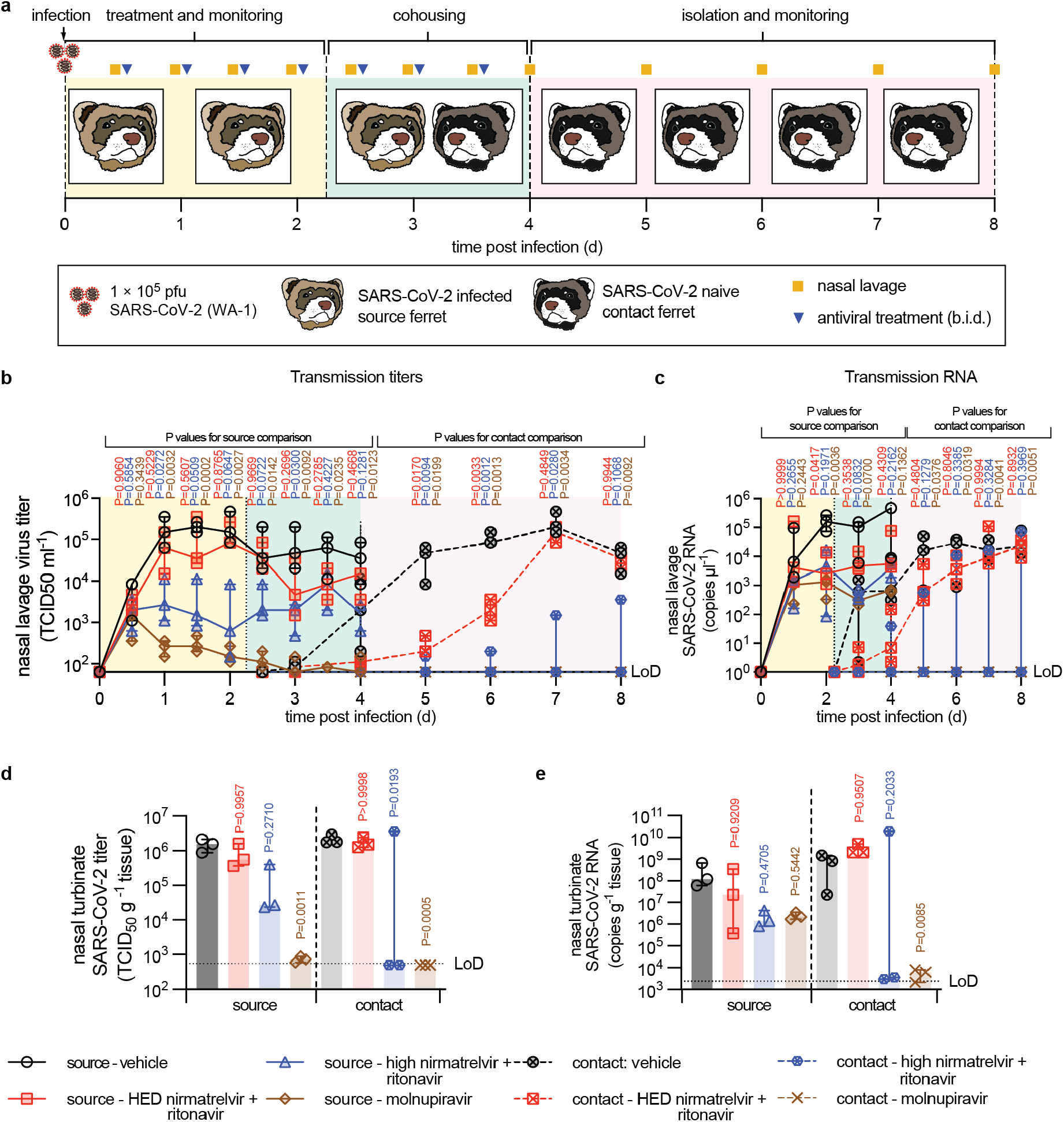
Nirmatrelvir does not block SARS-CoV-2 transmission. **a**, Direct-contact transmission study schematic. **b-c**, Infectious titers (b) and RNA copies (c) of SARS-CoV-2 present in nasal lavages of source and contact ferrets. **d-e**, Nasal turbinate titers (d) and RNA copies (e) in nasal turbinates extracted from source and contact animals on study days 4 and 8, respectively. Symbols represent independent biological repeats (results from individual animals), lines (b-c) and columns (d-e) denote medians. Error bars in (d-e) represent 95% CIs. Statistical analysis with 1-way (d-e) or 2-way (b-c) ANOVA with Dunnett’s (b-e) post-hoc tests; P values are shown, LoD limit of detection.

### Paxlovid treatment does not prevent transmission from SARS-CoV-2-infected ferrets

To examine the effect of paxlovid treatment on SARS-CoV-2 spread, we subjected treated ferrets to a direct-contact transmission study, first co-housing infected and treated source ferrets with untreated contact animals 42 hours after treatment initiation (Fig. 3a). Source animals were infected intranasally with SARS-CoV-2 WA1 as before, followed by paxlovid treatment in a b.i.d. regimen at HED or high-dose. For direct comparison with the only drug in clinical use that reportedly suppresses viral transmission in this model^23,24^, we included a molnupiravir group, treated orally b.i.d. with 5 mg/kg drug. All treatments were initiated 12 hours after infection. Equivalent amounts of shed infectious viral particles were detected in all source animal groups at the time of treatment start, confirming productive infection of all animals (Fig. 3b). None of the animals developed clinical signs (supplementary Fig. S2). Shed virus and RNA copy numbers of vehicle and paxlovid-treated sources resembled that of the efficacy study, whereas shed infectious titers in the molnupiravir arm rapidly declined, reaching detection level approximately 48 hours after treatment start (Fig. 3b-c).

SARS-CoV-2 spread rapidly to sentinels of vehicle-treated source animals. As noted before^23,24^, molnupiravir treatment of source animals was rapidly sterilizing, suppressing all transmission to untreated naïve contacts. By contrast, viral spread to sentinels of ferrets receiving HED paxlovid was delayed, but ultimately all contacts were infected and viral titers in nasal lavages were virtually identical to those of sentinels of vehicle-treated animals, indicating uninterrupted transmission. High-dose paxlovid reduced transmission rate, but productive viral transfer was still noted in one of the three contact pairs examined (Fig. 3b-c). Again, virus populations in lavage and turbinate samples of these animals had in most cases acquired mustelid-characteristic species adaptation mutations in the spike protein (extended data Fig. 3), but whole genome sequencing did not reveal any allele-dominant resistance mutation to nirmatrelvir in nsp5 (uploaded as NCBI BioProject PRJNA894555).

Viral titers and RNA copies in nasal turbinates determined 4 days after infection in source animals and 4 days after the end of co-housing in contact animals (study day 8) confirmed sterilizing antiviral activity of molnupiravir and complete suppression of all transmission to direct contacts (Fig. 3d-e). Sentinels of sources that had received HED paxlovid showed turbinate titers and viral RNA copies indistinguishable of those from animals that had been co-housed with vehicle-treated sources, and one of three contacts of high-dose paxlovid recipients showed vehicle-equivalent viral burden in upper respiratory tissues at study end. These results demonstrate that therapeutic treatment with HED paxlovid fails to interrupt viral transmission, and even at 5-times the human equivalent dose the drug only partially blocks SARS-CoV-2 spread.

### Prophylactic treatment of uninfected direct contacts with molnupiravir blocks infection

In the EPIC-PEP clinical trial, prophylactic administration of paxlovid to adults exposed to a household member with confirmed SARS-CoV-2 infection did not reduce the risk of infection^19,20^. Similar to the EPIC-PEP design, we prophylactically treated uninfected ferrets with HDE paxlovid or molnupiravir as before, followed 12 hours later by co-housing with infected, but untreated, source animals (Fig. 4a). Treatment of the sentinels was continued b.i.d. for 5.5 days, shed virus in nasal lavages of source and sentinels monitored daily, and virus load in the upper respiratory tract determined 6 days after infection.

**Figure 4.**
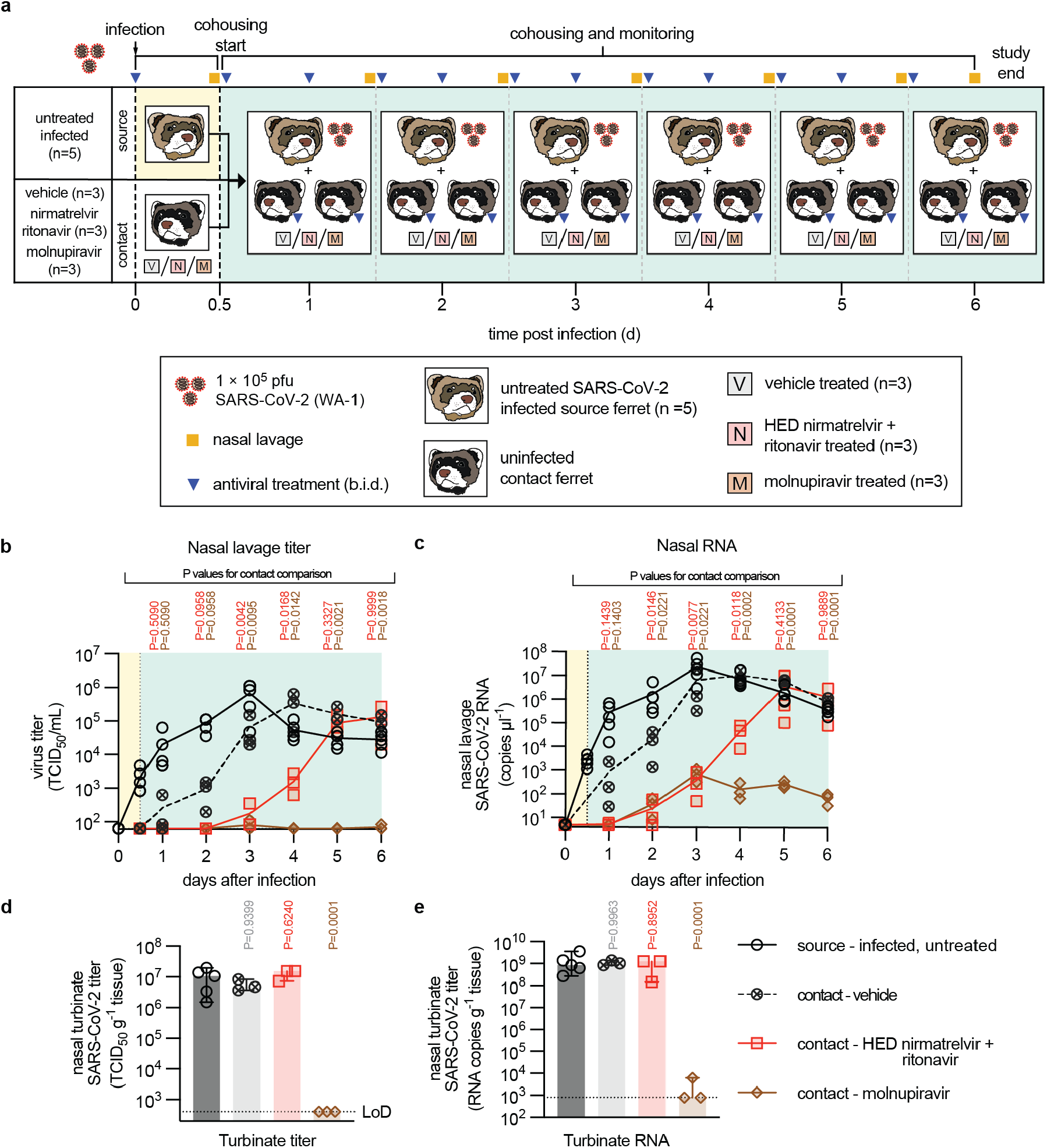
Prophylactic treatment with molnupiravir prevents productive infection by close contacts. **a**. Prophylaxis study schematic. **b-c**. Infectious titers (b) and RNA copies (c) of SARS-CoV-2 present in nasal lavages of source and contact ferrets. **d-e**, Nasal turbinate titers (d) and RNA copies (e) in nasal turbinates extracted from source and contact animals on study day 6. Symbols represent independent biological repeats (results from individual animals), lines (b-c) and columns (d-e) denote medians. Error bars in (d-e) represent 95% CIs. Statistical analysis with 1-way (d-e) or 2-way (b-c) ANOVA with Dunnett’s (b-e) post-hoc tests; P values are shown, LoD limit of detection.

We detected shed virus in lavages of all contacts receiving vehicle within 24 hours of co-housing, indicating productive transmission (Fig. 4b). Again none of the animals developed clinical signs (supplementary Fig. S3). HDE prophylactic paxlovid delayed transmission by approximately 48 hours, but ultimately all sentinels that received paxlovid were productively infected and shed virus titers reached by study end were indistinguishable from those of the vehicle group. In contrast, no infectious SARS-CoV-2 particles emerged in lavages of sentinels prophylactically treated with molnupiravir. Quantitation of viral RNA in the lavage samples largely mirrored the virus titer profiles (Fig. 4c). However, low amounts (<10^3^ RNA copies/µl) of SARS-CoV-2 RNA were present also in lavages of the molnupiravir group, demonstrating that co-housing with untreated source animals efficiently exposed sentinels of all three study arms to SARS-CoV-2. Nasal turbinates of all sentinels receiving molnupiravir did not contain infectious SARS-CoV-2 at study end, whereas virus burden in turbinates of animals treated with paxlovid was statistically indistinguishable from that found in the vehicle group (Fig. 4d). Viral RNA copies in turbinates mirrored the virus titer profiles, revealing high RNA copy numbers in all paxlovid recipients that were statistically equivalent to those derived from vehicle-treated animals (Fig. 4e). We found low RNA copies in the turbinates of one animal in the molnupiravir group, the other molnupiravir recipients remained viral RNA free.

These results demonstrate that prophylactic administration of paxlovid delays, but ultimately does not prevent, productive infection with SARS-CoV-2 from a direct-contact source, in essence recapitulating the outcome of the EPIC-PEP household transmission trial in the ferret model. In contrast, prophylactic molnupiravir completely suppressed productive infection, even when sentinels were exposed for several days to large amounts of SARS-CoV-2 shed from untreated source animals.

## Discussion

Paxlovid and molnupiravir were the first orally available antivirals that have received emergency authorization for the treatment of COVID-19 in the United States. Whereas initial clinical trials during VOC delta spread claimed major disparities in antiviral efficacy^12,13^, such a difference was not detected in large, retrospective analyses of drug performance during the VOC omicron surge. The Clalit study demonstrated that both paxlovid^16^ and molnupiravir^18^ significantly reduced hospitalizations and deaths (paxlovid hazard ratio (HR) 0.21, molnupiravir HR 0.26) due to COVID-19 in patients 65 years of age and older, who are at elevated risk of severe disease. Neither drug provided therapeutic benefit to younger adults. A retrospective cohort study from the Hong Kong VOC omicron wave returned similar results^17^; death rates were significantly reduced in recipients of either drug, patients experienced a lower risk of disease progression (paxlovid HR 0.57, molnupiravir HR 0.6), and the time to achieving a low viral burden after treatment start was significantly shortened (HR 1.38 for either drug). However, the study did not support a direct comparison of drug efficacy due to imbalances in baseline characteristics of the study groups such as patient comorbidities, age, and vaccine status^44^.

To assess efficacy of both drugs against COVID-19 of different severity under equivalent experimental conditions, we employed two relevant animal models of SARS-CoV-2 infection, the Roborovski dwarf hamster model for vulnerable patients at elevated risk of acute respiratory failure^24^ and the ferret upper respiratory tract disease and transmission model^23^. Single oral dose PK assessment in either model species ensured that dose levels applied in the study corresponded to nirmatrelvir plasma exposure equivalent to, or greater, that reached in human patients^31,32^, supporting physiological relevance of the assessment. Specifically, nirmatrelvir plasma exposure in the dwarf hamsters exceeded by over 7-times that seen in humans^31^, and administration of a calculated HED to ferrets resulted in plasma exposure virtually identical to that in humans, indicating good cross-species PK consistency.

In the dwarf hamster model of complicated COVID-19 with acute respiratory failure, assessment of molnupiravir and paxlovid efficacy supported three major conclusions: i) both drugs met the primary efficacy endpoints, since administration at HED prevented death of all treated animals; ii) both drugs statistically significantly reduced lung virus burden compared to vehicle-treated animals, and median effect size of virus titer reduction achieved by molnupiravir versus paxlovid was statistically equivalent; and iii) treatment benefit provided by either drug showed variations between individual animals when dwarf hamsters were infected with VOC omicron. We have reported recently that in the case of molnupiravir, biological sex of the treated dwarf hamsters contributes to effect size^24^. Although not powered for a full statistical analysis, paxlovid performance in the dwarf hamster model did not suggest an equivalent role of biological sex on effect size against VOC omicron. We favor the hypothesis that differences in the kinetics of viral host invasion between individual dwarf hamsters may contribute to individual variation of effect size after treatment of VOC omicron infection with paxlovid. *In toto*, the dwarf hamster model recapitulated the main conclusions of retrospective clinical studies in older adults of elevated risk to advance to severe COVID-19^17^; both drugs alleviated severe disease with comparable potency, preventing acute lung injury and death.

Prolonged self-isolation periods after a positive SARS-CoV-2 test impose considerable social hardship on the individual patient and economic burden on societies^45-47^, which has resulted in eroding public compliance with self-quarantine during the course of the pandemic^48^. In addition to direct therapeutic benefit, pharmacologically shortening the duration of the infectious period will improve patient quality of life and may provide epidemiologic benefit through interruption of viral transmission chains^49,50^. We have established efficient suppression of SARS-CoV-2 transmission by molnupiravir^23,24^ and experimental nucleoside analog antivirals^33^ in previous work. Considering the comparable efficacy of paxlovid and molnupiravir to control virus replication in the lower respiratory tract, we were surprised that HED paxlovid treatment, in contrast to molnupiravir, was insufficient to eliminate upper respiratory tract virus burden and failed to prevent SARS-CoV-2 transmission in ferrets, independent of whether source animals received paxlovid therapeutically or direct contacts were treated prophylactically. Whereas therapeutic molnupiravir had ultimately a sterilizing effect in infected ferrets and prophylactic molnupiravir prevented productive infection, shed virus titers in animals that received therapeutic or prophylactic paxlovid plateaued and/or increased towards the end of the treatment course. Two lines of evidence, clinical reports of rebounding virus replication in human patients who have received paxlovid^2,21,22^ and failure of the EPIC-PEP trial of prophylactic use of paxlovid^19,20^, underscore relevance of these ferret-derived data for human disease. If the model predicts the impact of molnupiravir on virus transmission with equal accuracy, prophylactic use of molnupiravir may have the potential to lower the risk of close contact transmission.

Despite extensive resistance profiling attempts^10,40,51^, no viral escape from molnupiravir has yet been reported, whereas pe-existing paxlovid resistance mutations have been detected in some circulating SARS-CoV-2 strains that are considered able to spread^42,52^. However, whole genome sequencing of virus population isolated from paxlovid-experienced ferrets provided no evidence that replication rebound and transmission both from and to paxlovid-treated animals in our study were supported by emerging resistance to nirmatrelvir^41-43^. Alternatively, poor control of upper respiratory infection and failure to prevent transmission could be due to lower nirmatrelvir exposure in upper respiratory tissues than lung or changing upper respiratory exposure profiles after repeat dosing. In addition, signature mustelid adaptation mutations in the spike protein became allele-dominant in most infected ferrets, suggesting a combination of species adaptation of spike and low nirmatrelvir exposure in privileged upper respiratory compartments as a possible cause for viral rebound and transmission in paxlovid-treated ferrets.

This study affirms retrospective clinical analyses that early treatment of patients at elevated risk of progression to severe COVID-19 with either paxlovid or molnupiravir will provide significant therapeutic benefit. In contrast to efficient suppression of SARS-CoV-2 transmission by molnupiravir, however, paxlovid had only a moderate impact on controlling virus replication in the upper respiratory tract and did not shorten the infectious period in ferrets or lower the risk of infection when administered prophylactically. Considering reports of virus rebound in human patients after paxlovid that phenocopy these ferret data, paxlovid may not be suitable to provide epidemiologic benefit by interrupting SARS-CoV-2 transmission chains.

## Methods

### Ethics statement

All *in vivo* studies were performed in compliance with the Guide for the Care and Use of Laboratory Animals, National Institutes of Health guidelines, and the Animal Welfare Act Code of Federal Regulations. Experiments with SARS-CoV-2 involving hamsters and ferrets were approved by the Georgia State Institutional Animal Care and Use Committee under protocol A21019 and A20031, respectively. All experiments involving infectious SARS-CoV-2 were performed in BSL-3/ABSL-3 facilities at the Georgia State University, and were approved by the Georgia State Institutional Biosafety Committee under protocol B20016.

### Study Design

Roborovski dwarf hamsters (*Phodopus roborovskii*, 3-4 months old) and Ferrets (*Mustela putorius furo*, 6-10 months old) were used as *in vivo* models to assess the therapeutic efficacy of orally administered nirmatrelvir against infections with SARS-CoV-2. Prior to studies, hamsters and ferrets were rested for at least five days. Roborovski dwarf hamsters were used to study the effects of nirmatrelvir treatment on severe disease associated with lower respiratory tract infection, acute lung injury, and death. Ferrets were used the examine the effect of nirmatrelvir on upper respiratory infection and transmission. SARS-CoV-2 infections were administered to animals through intranasal inoculation. For hamsters, animals were monitored regularly for clinical signs and viral loads were determined in respiratory tract tissues at endpoint. For ferrets, upper respiratory tract viral titers were assessed at least once daily through nasal lavages. In addition, upper respiratory tract tissues were harvested at endpoint to assess upper respiratory tract viral loads. Virus loads were determined by plaque assays or TCID_50_ and RT-qPCR quantitation.

### Experimental details

Detailed description of experimental procedures and reagents specifying cells and viruses used, virus yield reduction assays, SARS-CoV-2 titration by plaque assay, Single dose pharmacokinetics in hamsters, hamster efficacy studies, single ascending dose pharmacokinetics in ferrets, *in vivo* efficacy testing of nirmatrelvir in ferrets against VOCs, the effect of nirmatrelvir treatment on SARS-CoV-2 contact transmission in ferrets, titration of SARS-CoV-2 in tissue extracts, quantitation of SARS-CoV-2 RNA, and SARS-CoV-2 genome sequencing are provided as supplementary information.

### Statistics and reproducibility

The Microsoft Excel (versions 16.52) and Numbers (version 10.1) software packages were used for data collection. The GraphPad Prism (version 9.3.1) software package was used for data analysis. Reverse transcription RT-qPCR data were collected and analyzed using the QuantStudio Design and Analysis (version 1.5.2; Applied Biosystems) software package. Figures were assembled and generated with Adobe Illustrator (version CS6). T-tests were used to evaluate statistical significance between experiments with two sets of data. To evaluate statistical significance when more than two groups were compared or datasets contained two independent variables, 1- and 2-way ANOVAs, respectively, were used with Tukey’s or Dunnett’s multiple comparisons post-hoc tests as specified in figure legends. Specific statistical tests are specified in the figure legends for individual studies. Supplementary dataset 1 summarizes the statistical analyses (effect sized, P values, and degrees of freedom) of the respective datasets. Effect sizes between groups were calculated as η^2^= (SS_effect_) / (SS_total_) for one-way ANOVA and ω^2^= (SS_effect_ – (df_effect_)(MS_error_)) / MS_error_ + SS_total_ for two-way ANOVA; SS_effect_, sum of squares for the effect; SS_total_, sum of squares for total; df_effect_, degrees of freedom for the effect; MS_error_, mean square error. Effective concentrations for antiviral potency were calculated from dose-response virus yield reduction assay datasets through four-parameter variable slope regression modelling. A biological repeat refers to measurements taken from distinct samples, and the results obtained for each individual biological repeat are shown in the figures along with the exact size (n, number) of biologically independent samples, animals, or independent experiments. The measure of the center (connecting lines and columns) is specified in the figure legends. The statistical significance level (α) was set to <0.05 for all experiments. Exact P values are shown in the supplementary dataset 1 and in individual graphs when possible. All quantitative source data are provided in the supplementary dataset 2.

### Reporting summary

Further information on research design is available in the Nature Research Reporting Summary linked to this article.

### Data availability

The amplicon tiling sequencing reads generated in this study have been deposited in the NCBI BioProject database under accession code PRJNA894555. All other data generated in this study are provided in this published article and the Supplementary Information/Source Data file. Source data are provided in this paper.

### Code availability

Sequencing reads were analyzed using the TAYLOR pipeline (available at https://github.com/greninger-lab/covid_swift_pipeline). All commercial computer codes and algorithms used are specified in the Methods section.

## Author contributions

R.M.C., C.M.L., and R.K.P. conceived and designed the experiments. R.M.C., C.M.L., J.D.W., A.K., and R.K.P. conducted most of the experiments. R.M.C. created all figure schematics. A.L.G., N.A.P.L, P.R. performed next-generation sequencing. M.K.A., R.E.K., Z.M.S., and A.A.K. performed mass spectrometry analysis. M.G.N. and G.R.P. provided critical materials. R.M.C., C.M.L., N.A.P.L, A.L.G., and R.K.P. analyzed the data. R.M.C. and R.K.P. wrote the manuscript.

## Acknowledgements

We thank the Georgia State University Department of Animal Resources and High Containment Core for Assistance and J-J Yoon for technical support. This study was supported, in part, by public health service grants AI141222 (to R.K.P.) and AI171403 project 1 (to R.K.P.) and Scientific Core D (to A.L.G.) from the NIH/NIAID.

## Conflict of interest

GRP, MGN and AAK receive licensing fees and royalties based on Emory’s sublicense of the molnupiravir technology to Ridgeback Biotherapeutics, and a family member of GRP serves as the Chief Medical Officer of Ridgeback. This technology is the subject of the research described in this paper. The terms of this arrangement have been reviewed and approved by Emory University in accordance with its conflict-of-interest policies. RKP reports contract testing from Enanta Pharmaceuticals and Atea Pharmaceuticals, and research support from Gilead Sciences, outside of the described work. ALG reports contract testing from Abbott, Cepheid, Novavax, Pfizer, Janssen and Hologic and research support from Gilead Sciences and Merck, outside of the described work. All other authors declare that they have no competing interests to report.

## Extended data Tables and Figures

**Extended data Table 1.**
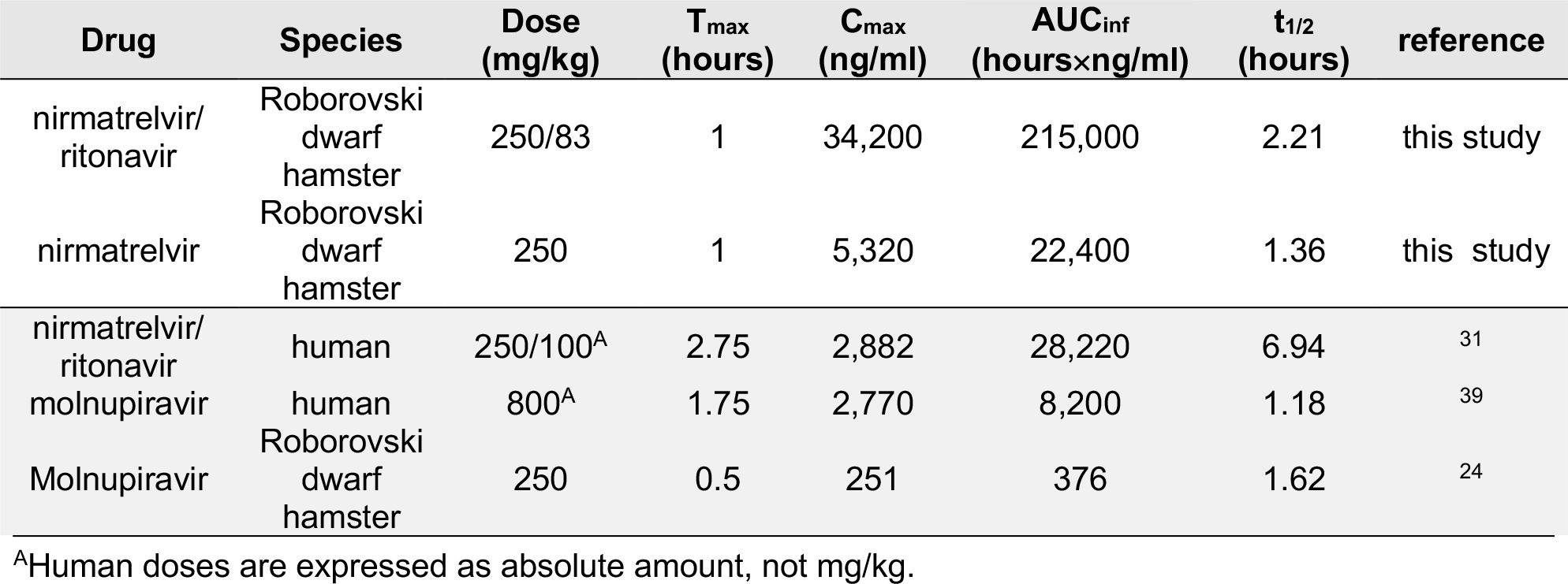
Single dose pharmacokinetics or nirmatrelvir in Roborovski dwarf hamster plasma.

**Extended data Table 2.** Single dose pharmacokinetics of nirmatrelvir in ferret plasma.

| Drug | Species | Dose (mg/kg) | T<sub>max</sub> (hours) | C<sub>max</sub> (ng/ml) | AUC<sub>inf</sub> (hours×ng/ml) | t<sub>1/2</sub> (hours) | reference |
| --- | --- | --- | --- | --- | --- | --- | --- |
| nirmatrelvir/<br>ritonavir | ferret | 20/6 | 2.33 | 8990 | 32700 | 1.05 | this study |
| nirmatrelvir/<br>ritonavir | ferret | 100/6 | 4 | 11800 | 49500 | 0.74 | this study |
| nirmatrelvir/<br>ritonavir | human | 250/100<sup>A</sup> | 6.94 | 2882 | 28220 | 6.94 | 31 |
| molnupiravir | human | 800<sup>A</sup> | 1.75 | 2770 | 8,200 | 1.18<br>(7.28) | 39 |
| molnupiravir | ferret | 4 | 1.7 | 907 | 3,421 | 8.2 | 40 |
| molnupiravir | ferret | 7 | 1.5 | 2851 | 7569 | 9.2 | 40 |
| molnupiravir | ferret | 20 | 1.8 | 3992 | 18,793 | 4.7 | 40 |
| molnupiravir | macaque | 130 | 1.62 | 2644 | 25,299 | 1.77 | 40 |

<sup>A</sup>Human doses are expressed as absolute amount, not mg/kg.

**Extended data Fig. 1.**
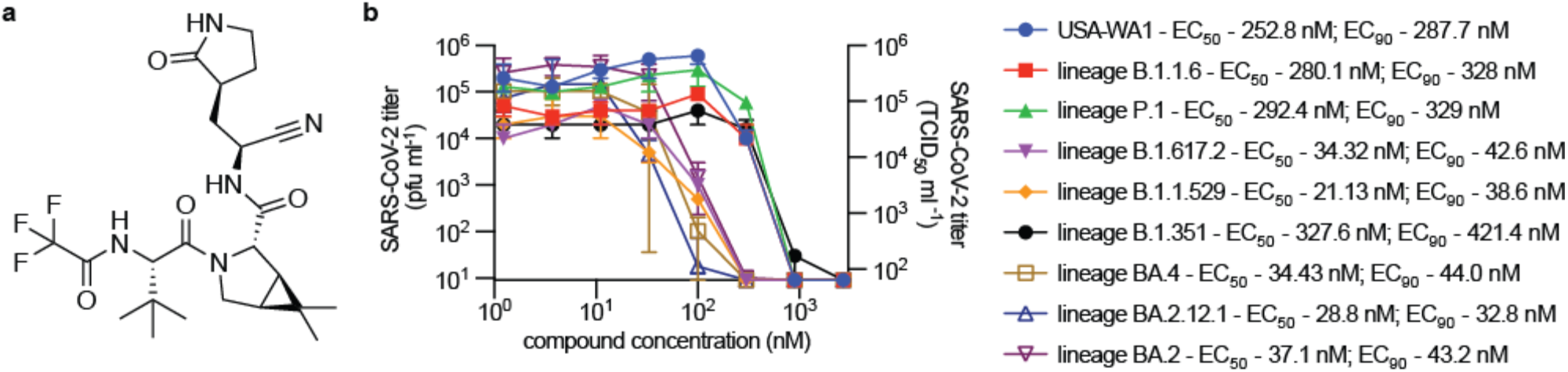
*In vitro* antiviral activity of nirmatrelvir. **a**, Structure of nirmatrelvir. **b**, Nirmatrelvir dose-response assays against SARS-CoV-2 WA1, and lineages B.1.1.7 (VOC α), B.1.351 (VOC β), P.1 (VOC γ), B.1.617.2 (VOC δ), B.1.1.529, BA.2, B1.2.12.1, and BA.4 on VeroE6-TMPRSS2 cells. Numbers denote 50 and 90% inhibitory concentrations (EC_50_ and EC_90_, respectively), determined through non-linear 4-parameter variable slope regression modeling. Lines intersect group medians; error bars represent 95% confidence intervals (CIs).

**Extended data Fig. 2.**
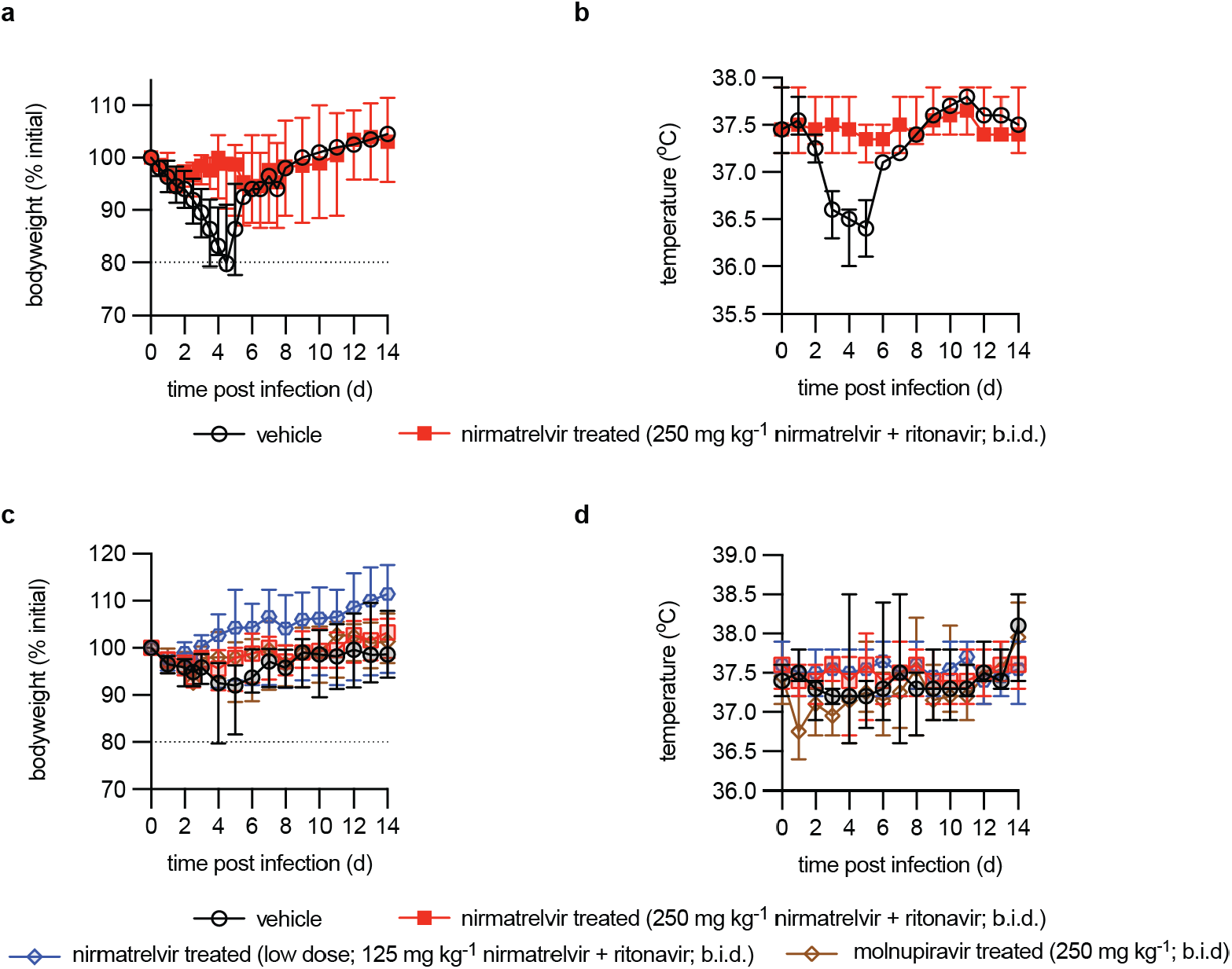
Clinical signs in Roborovski dwarf hamsters. **a-b**, Bodyweight (a) and temperature (b) of VOC delta infected dwarf hamsters. **c-d**, Bodyweight (c) and temperature (d) of VOC omicron infected dwarf hamsters. Symbols represent group medians, lines intersect medians, and error bars represent 95% CIs.

**Extended data Fig. 3.**
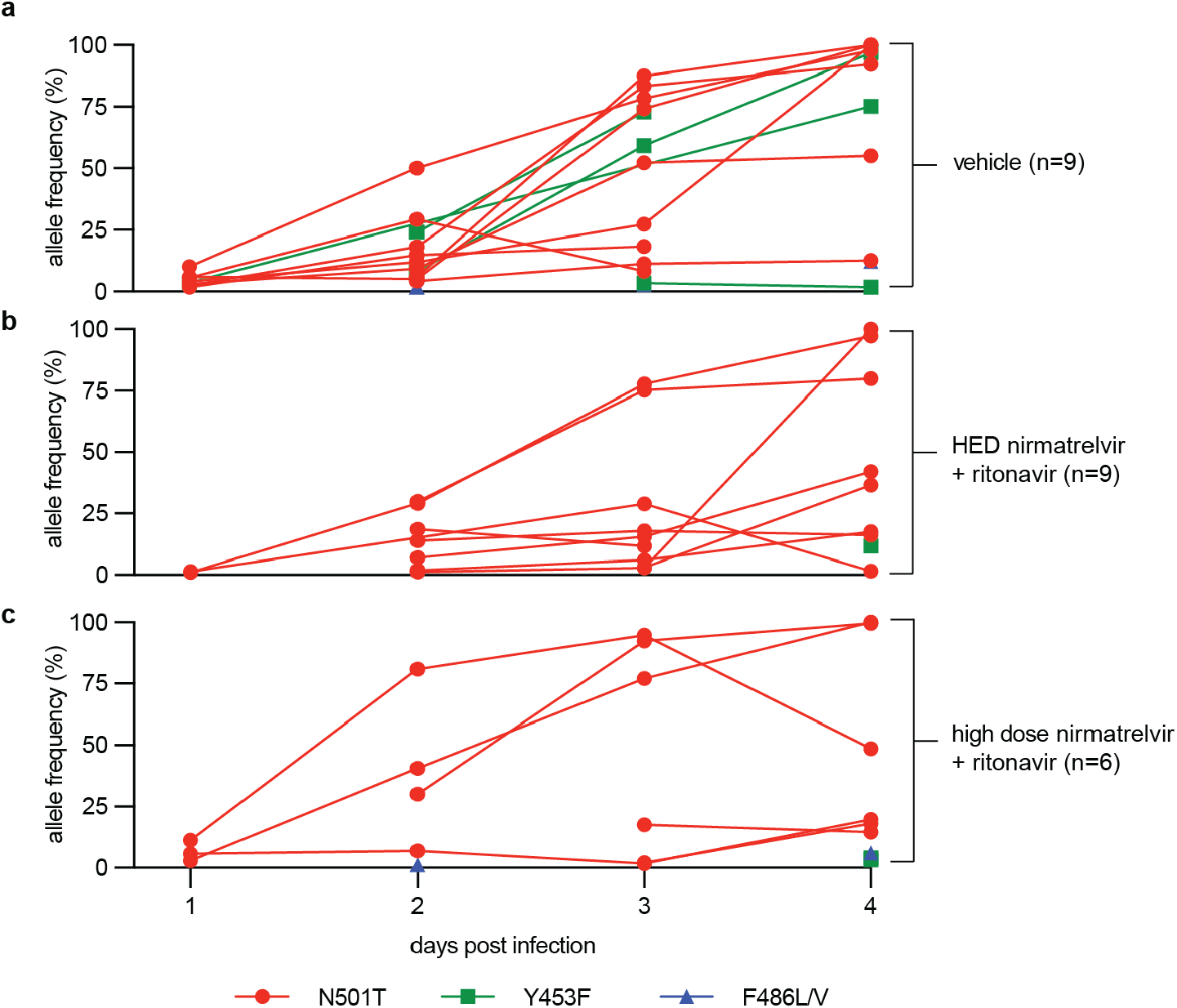
Genetic variation in virus populations recovered from treatment-experienced animals. **a-c**, Relative allele frequency of all mutations that emerged in the spike protein of SARS-CoV-2 populations in nasal lavage samples extracted from infected ferrets specified in Fig. 2b and Fig. 3a. Whole genome sequencing results are shown for animals treated with vehicle (a), HED paxlovid (b), or high dose paxlovid (c) and identified mutations N501T, Y453F, and F486L/V that are characteristic for SARS-C-V-2 adaptation to mustelids. Lines connect relative allele frequencies for independent biological repeats (individual animals) on days 1 to 4 after infection. On days 1-3 viral RNA was isolated from nasal lavage samples, day 4 represents viral RNA extracted from nasal turbinates.

## Notes

https://www.ncbi.nlm.nih.gov/bioproject/?term=PRJNA894555

### Competing Interest Statement

GRP, MGN and AAK receive licensing fees and royalties based on Emorys sublicense of the molnupiravir technology to Ridgeback Biotherapeutics, and a family member of GRP serves as the Chief Medical Officer of Ridgeback. This technology is the subject of the research described in this paper. The terms of this arrangement have been reviewed and approved by Emory University in accordance with its conflict-of-interest policies. RKP reports contract testing from Enanta Pharmaceuticals and Atea Pharmaceuticals, and research support from Gilead Sciences, outside of the described work. ALG reports contract testing from Abbott, Cepheid, Novavax, Pfizer, Janssen and Hologic and research support from Gilead Sciences and Merck, outside of the described work. All other authors declare that they have no competing interests to report.

