## Supplementary Information for "Paxlovid-like nirmatrelvir/ritonavir fails to block SARS-CoV-2 transmission in ferrets"

### Experimental reagents and methods

#### Cells

VeroE6-TMPRSS2 (BPS Bioscience #78081) and Calu-3 (ATCC HB-55™) cells were cultivated at 37°C with 5% $CO_2$ in Dulbecco's Modified Eagle's Medium (DMEM) supplemented with 7.5% heat-inactivated fetal bovine serum (FBS). All cells were authenticated and checked routinely for mycoplasma prior to use.

#### Viruses

SARS-CoV-2 USA-WA1 (lineage A, isolate USA-WA1/2020, BEI cat# NR-52281) was obtained from BEI resources and amplified on Calu-3 cells. SARS-CoV-2 isolates for lineage B.1.1.7 (VOC α; hCoV-19/USA/CA/UCSD_5574/2020), lineage B.1.351 (VOC β; hCoV-19/South Africa/KRISP-K005325/2020), lineage P.1 (VOC γ; hCoV-19/Japan/TY7-503/2021), lineage B.1.617.2 (VOC δ; clinical isolate #233067), lineage B.1.1.529 (VOC o; hCoV-19/USA/WA-UW-21120120771/2021) lineage BA.2 (clinical isolate 22012361822A), lineage BA.2.12.1 (hCoV-19/USA/WA-CDC-UW22050170242/2022), and lineage BA.4 (hCoV-19/USA/WA-CDC-UW22051283052/2022) were obtained from UW Virology and amplified on Calu-3 cells. All viruses were authenticated by whole genome next generation sequencing prior to use.

#### Virus yield reduction

Nirmatrelvir was dissolved in water-free DMSO at 10 mM stock concentration and stored in single-use aliquots at -80°C. Cells were seeded 14 hours prior to experimentation in 24-well plates at $2 \times 10^5$ cells per well in 400 µl per well DMEM + 7.5% FBS supplemented with 2 µM CP-100356. Serial dilutions were generated in culture media containing 2 µM CP-100356 spanning a final concentration range from 2700 to 1.4 nM (3-fold serial dilution; 8 steps; dilution series were prepared 2-fold overconcentrated (54000 to 2.8 nM)). Every concentration of each test article was assessed in 3 biological (independent) repeats. Vehicle-treated control cultures DMSO volume equivalents caorresponding to the highest test compound concentration. Serial dilutions were transferred to target wells (500 µl per well), immediately followed by infection with SARS-CoV-2 (multiplicity of infection 0.1 pfu per cell) in 100 µl per well (viral dilution media contained 2 µM CP-100356). Infected plates were incubated in a humidified incubator at 37°C and 5% $CO_2$ for 48 hours. Progeny virus titers in culture supernatants were determined through plaque-forming or $TCID_{50}$ assay on Vero-E6-TMPRSS2 cells. Inhibitory concentrations were calculated based on 4-parameter variable slope regression models.

**SARS-CoV-2 titration by plaque assay**

All samples were serially diluted in DMEM media supplemented with 2% FBS and antibiotic-antimycotic solution (Gibco). Dilutions were added to VeroE6-TMPRSS2 cells seeded 24 hours earlier in 12-well plates at $3 \times 10^5$ cells per well. Dilutions were allowed to adsorb for 2 hours at 37°C. Following adsorption, inoculum was removed, and cells were overlaid with 1.2% Avicel (FMC BioPolymer) in DMEM supplemented with antibiotic-antimycotic solution and incubated at 37°C with 5% $CO_2$. After incubating for 3 days, Avicel was removed, cells were washed with PBS and fixed with 10% neutral buffered formalin. Plaques were visualized using 1% crystal violet.

**Single dose pharmacokinetics in hamsters**

Groups of hamsters were administered a single oral dose of nirmatrelvir (250 mg/kg; 0.5% MC with 2% Tween80) with or without ritonavir (83.4 mg/kg; 20% ethanol). Blood was harvested in $K_2$EDTA capillary tubes (Sarsdet) 0.5, 1, 2, 4, 8 and 24 hours after dosing. Plasma was clarified by centrifugation (4°C, 2000 rpm, 5 min). To determine tissue concentrations of nirmatrelvir, groups of hamsters were sacrificed 1, 8 and 24 hours after dosing and selected tissues were harvested and stored at -80°C.

**Hamster efficacy studies**

Groups of male and female Roborovski dwarf hamsters were anesthetized using ketamine/dexmedetomidine and infected with $10^4$ pfu per animal (50 µl total volume, 25 µl per nare). Anesthesia reversed with atipamezole. Hamsters were monitored daily (bodyweight, temperature, and clinical score). Treatment started 12 hours after infection with nirmatrelvir (200 µl, 0.5% MC with 2% Tween80; 250 mg/kg or 125 mg/kg b.i.d.) and ritonavir (50 µl, 20% EtOH; 83.3 mg/kg, b.i.d.). Treatment continued twice daily for 5 and 7 days after infection for VOC δ and VOC o, respectively. Treatment was discontinued for VOC δ after 5 days due to treated hamsters developing pronounced diarrhea. Groups of hamsters were euthanized three days after infection to assess lower respiratory tract viral load. Lungs were harvested, washed with sterile phosphate buffered saline, and homogenized in PBS supplemented with antibiotic-antimycotic using a bead blaster (3 bursts of 30 seconds at 4°C each, 30 seconds pause between bursts). Lung homogenates were clarified by centrifugation (20,000 × g, 10 minutes, 4°C), aliquoted and stored at -80°C until processed. For RNA quantification, lungs and tracheas were harvested and homogenized in RNeasy RNA lysis buffer (Qiagen) in accordance with the manufacturers protocol.

**Single ascending dose pharmacokinetics in ferrets**

Groups of ferrets were administered a single oral dose of nirmatrelvir at a dosage of 20 mg/kg or 100 mg/kg; (2 ml; 0.5% MC with 2% Tween80) with or without ritonavir (1 ml; 6 mg/kg; 20% ethanol). A dosage of 6 mg/kg ritonavir was selected based on previous studies in ferrets (*53*). Blood was harvested in $K_2$EDTA capillary tubes (Sarstedt Microvette CB300) at 0.25, 0.5, 1, 2, 4, 6, 8 and 12 hours after dosing. Plasma was clarified by centrifugation (4°C, 2,000 × g, 5 min), aliquoted and stored at -80°C. Nirmatrelvir concentrations were analyzed using a qualified LC/MS/MS method, and PK parameters were calculated using non-compartmental analysis with WinNonlin 8.3.3.33.

***In vivo* efficacy of nirmatrelvir in ferrets against VOCs**

Groups of ferrets (n=3-6) were inoculated with $1 \times 10^5$ pfu of SARS-CoV-2 USA-WA1 (lineage A, isolate USA-WA1/2020, BEI cat# NR-52281; 1 ml, 0.5 ml per nare). Treatment was initiated 12 hours after infection and continued twice daily until study end. Ferrets were treated with vehicle (0.5% MC with 2% Tween80), nirmatrelvir-ritonavir (20 or 100 mg/kg nirmatrelvir; 6 mg/kg ritonavir). Treatments were administered by oral gavage 12 hours until 4 days after infection. Nasal lavages were collected every 12 h after study start. All ferrets were euthanized 4 days after infection and tissues were harvested to determine SARS-CoV-2 titers and the presence of viral RNA.

**Effect of nirmatrelvir treatment on SARS-CoV-2 contact transmission in ferrets**

A group of source ferrets (n=12) were inoculated intranasally with $1 \times 10^5$ pfu of SARS-CoV-2 USA-WA1 (lineage A, isolate USA-WA1/2020, BEI cat# NR-52281; 1 ml, 0.5 ml per nare). Twelve hours after infections, source ferrets were further divided into four groups (n=3), receiving vehicle, molnupiravir (5 mg/kg, b.i.d.), nirmatrelvir plus ritonavir at a dose of either 20 or 100 mg/k nirmatrelvir with 6 mg/k ritonavir administered by oral gavage. At 54 h after infection, each source ferret was co-housed with one uninfected and untreated contact ferret. Cohousing continued until 96 hours after infection, when sourced ferrets were euthanized and contact ferrets were separated and housed individually. All contact ferrets were monitored for 4 days after separation from source ferrets and then euthanized. Nasal lavages were performed on source ferrets twice daily until cohousing ended. After cohousing started, nasal lavages were performed on all contact ferrets every 24 hours. Upon euthanizing ferrets, nasal turbinates were harvested to determine infectious titers and the presence of viral RNA.

**Titration of SARS-CoV-2 in tissue extracts**

For virus titration, the organs were weighed and homogenized in PBS supplemented with antibiotic-antimycotic solution. Tissues were homogenized using a bead blaster (3 bursts of 30 seconds at 4°C each, 30 seconds pause between bursts). Tissue homogenates were clarified by centrifugation (20,000 × g for 5 min at 4 °C). The clarified supernatants were harvested, frozen and used in subsequent $TCID_{50}$ or plaque assays. For detection of viral RNA, the harvested organs were homogenized in lysis buffer (Qiagen), and the total RNA was extracted using a RNeasy mini kit (Qiagen). For nasal lavages, RNA was extracted using a Quick-RNA Viral Kit (Zymo) in accordance with the manufacturer's protocols.

**Quantitation of SARS-CoV-2 RNA**

SARS-CoV-2 RNA was detected using the nCoV_IP2 primer-probe set (nCoV_IP2-12669Fw ATGAGCTTAGTCCTGTTG; nCoV_IP2-12759Rv CTCCCTTTGTTGTGTTGT; nCoV_IP2-12696bProbe(+) 5'FAM-AGATGTCTTGTGCTGCCGGTA-3'BHQ-1) (National Reference Center for Respiratory Viruses, Institut Pasteur). RT-qPCR reactions were performed using an QuantStudio 3 real-time PCR system using the QuantStudio Design and Analysis package. The nCoV_IP2 primer-probe set was used with Taqman Fast Virus 1-step master mix (Thermo Fisher Scientific) to detect viral RNA. A PCR fragment (nt 12669-14146 of the SARS-CoV-2 genome) generated from viral complementary DNA using the nCoV_IP2 forward primer and the nCoV_IP4 reverse primer (nCoV_IP4-14146Rv CTGGTCAAGGTTAATATAGG) was used to create a standard curve to quantitate the RNA copy numbers. RNA copy numbers were normalized to the weight of tissues used.

**SARS-CoV-2 genome sequencing**

SARS-CoV-2 positive specimens were sequenced using IDT xGen SARS-CoV-2 Amplicon Panel (previously Swift Biosciences SNAP panel) (*54*), following the manufacturer's protocol. Sequencing reads were processed and analyzed using TAYLOR default settings (*54, 55*) available at https://github.com/greninger-lab/covid_swift_pipeline. Sequencing reads are available in NCBI BioProject PRJNA894555.

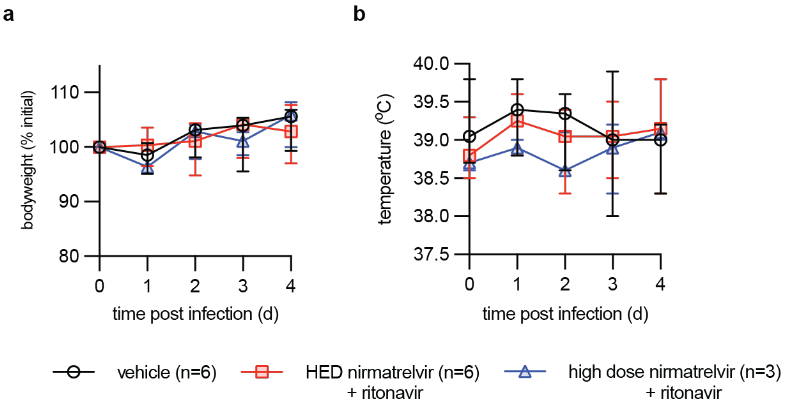


**Supplementary Fig. S1. Clinical signs of SARS-CoV-2 infected ferrets in efficacy study. a-b**, Bodyweight (a) and temperature (b) of SARS-CoV-2 infected ferrets from figure 2. Symbols represent group medians, lines intersect medians, and error bars represent 95% CIs.

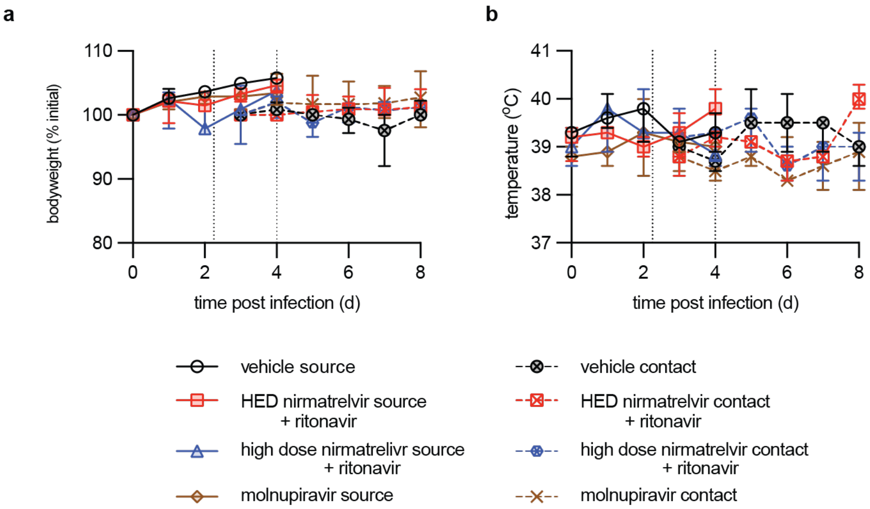


**Supplementary Fig. S2. Clinical signs of SARS-CoV-2 infected ferrets in the transmission study. a-b**, Body weight (a) and temperature (b) of SARS-CoV-2 infected ferrets from figure 3. Symbols represent group medians, lines intersect medians, and error bars represent the 95% CIs. Dashed vertical lines represent the time period of co-housing of source and naïve contact ferrets.

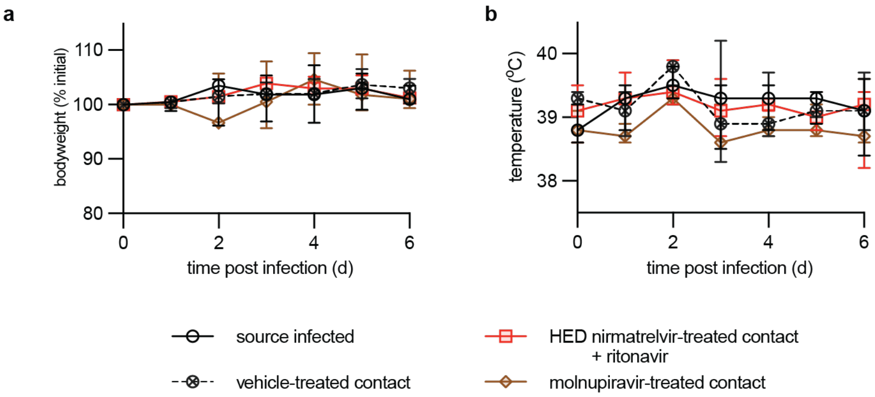

**Supplementary Fig. S3. Clinical signs of SARS-CoV-2 infected ferrets in the prophylactic treatment study. a-b**, Body weight (a) and temperature (b) of SARS-CoV-2 infected ferrets from figure 4. Symbols represent group medians, lines intersect medians, and error bars represent the 95% CIs. Infected source and prophylactically treated contact ferrets were co-housed starting 12 hours after infection.
